# Indirect Vibration of the Upper Limbs Alters Transmission Along Spinal but not Corticospinal Pathways

**DOI:** 10.1101/2020.10.15.341040

**Authors:** Trevor S. Barss, David F. Collins, Dylan Miller, Amit N. Pujari

## Abstract

The aim of this study was to investigate whether indirect upper limb vibration (ULV) modulates transmission along spinal and corticospinal pathways that control the human forearm. All measures were assessed under CONTROL (no vibration) and ULV (30 Hz; 0.4 mm displacement) conditions while participants maintained a small contraction of the right flexor carpi radialis (FCR) muscle. To assess spinal pathways, Hoffmann reflexes (H-reflexes) elicited by stimulation of the median nerve were recorded from FCR with motor response (M-wave) amplitudes matched between conditions. An H-reflex conditioning paradigm was also used to assess changes in presynaptic inhibition by stimulating the superficial radial (SR) nerve (5 pulses at 300Hz) 37 ms prior to median nerve stimulation. Cutaneous reflexes in FCR elicited by stimulation of the SR nerve at the wrist were also recorded. To assess corticospinal pathways, motor evoked potentials (MEPs) elicited by transcranial magnetic stimulation of the contralateral motor cortex were recorded from the right FCR and biceps brachii (BB). ULV significantly reduced H-reflex amplitude by 15.7% for both conditioned and unconditioned reflexes (24.0±15.7 vs 18.4±11.2 % M_max_; *p*<0.05). Middle latency cutaneous reflexes were also significantly reduced by 20.0% from CONTROL (−1.50 ± 2.1 % Mmax) to ULV (−1.73 ± 2.2 % Mmax; *p*<0.05). There was no significant effect of ULV on MEP amplitude (*p*>0.05). Therefore, ULV inhibits cutaneous and H-reflex transmission without influencing corticospinal excitability of the forearm flexors suggesting increased presynaptic inhibition of afferent transmission as a likely mechanism. A general increase in inhibition of spinal pathways with ULV may have important implications for improving rehabilitation for individuals with spasticity (SCI, stroke, MS, etc).

## 1 Introduction

The use of vibration during exercise and rehabilitation continues to gain popularity as a modality to improve function and performance (Cochrane, 2011; Lai et al., 2018). When vibration is used in this context it can be broadly classified into two categories, 1) stimulation directly applied to a specific muscle or tendon, and 2) indirect vibration which is not muscle specific and is delivered either through the feet while standing on a platform or through the hands by holding a device. Indirect vibration delivered through the hands is commonly referred to as upper limb vibration (ULV) while indirect vibration delivered to the lower limbs is referred to as whole-body vibration (WBV). Vibration applied directly to a muscle or tendon has a long history within the literature (Hagbarth KE, 1965). More recently, indirect vibration has been investigated as a potential assistive modality in both neurologically intact and neurologically impaired individuals (Marín et al., 2010).

Most indirect vibration research has focused on WBV, which in neurologically intact individuals, can improve muscle strength (Marín et al., 2010), muscle power (Marín and Rhea, 2010), fall risk (Jepsen et al., 2017), flexibility (Houston et al., 2015), and balance (Tseng et al., 2016). However, the effect of WBV on bone mineral density (Dionello et al., 2016) or lean mass (Chen et al., 2017; Lai et al., 2018) are less clear. For example, a recent meta-analysis indicated WBV may lead to improvements in lean muscle mass within younger adults but had no influence in children, adolescents, postmenopausal women, or an aging population (Chen et al., 2017). Within clinical populations, the results have also been equivocal. For example, WBV reduced muscle spasticity and improved balance and walking for individuals with an incomplete spinal cord injury (SCI) (In et al., 2018) however, a recent systematic review and meta-analysis indicates that WBV had no beneficial effect on muscle strength, balance and gait performance for individuals experiencing a chronic stroke (Lu et al., 2015). Similarly, WBV did not improve functional performance for individuals with neurodegenerative diseases (Parkinson’s or multiple sclerosis) compared to other active physical therapy or passive interventions (Sitjà Rabert et al., 2012).

While less research has focused on ULV, there is some evidence that it has potential to be an effective exercise and rehabilitation strategy. In neurologically intact participants, ULV increased the rate of force development during maximal isometric elbow flexion (de Paula et al., 2017) and ULV improved mobility and motor function for individuals with upper limb hemiparesis after a stroke (Oliveira et al., 2018). As well, ULV improved grip strength and shoulder range of motion compared to a control group in breast cancer patients (Kneis et al., 2018). Unfortunately, a lack of knowledge of the pathways and mechanisms being altered during ULV continues to limit its effective implementation (Cochrane, 2011).

Direct vibration of a muscle or tendon alters transmission of primary and secondary muscle afferents (Ia, Ib, and type II afferents) (Eklund and Hagbarth, 1966; Bishop B., 1974; Burke et al., 1976), cutaneous mechanoreceptors (Freeman and Johnson, 1982), and modulates cortical excitability (Münte et al., 1996). The effects of indirect vibration on spinal and corticospinal pathways, however, have yet to be clearly established. In the lower limbs of neurologically intact individuals, WBV inhibits both stretch reflexes (Ritzmann et al., 2013) and H-reflexes(Armstrong et al., 2008; Kipp et al., 2011; Ritzmann et al., 2013; Hortobágyi et al., 2014; Ahmadi et al., 2015; Harwood et al., 2017; Laudani et al., 2018) and increases disynaptic reciprocal inhibition (Ritzmann et al., 2018). WBV also inhibits H-reflexes in lower limb muscles of individuals with a SCI (Sayenko et al., 2010). In summary, the diminished H-reflex and stretch reflex amplitudes indicate attenuated sensorimotor transmission at the level of the spinal cord of either a pre- or post-synaptic nature during and after WBV exercise. WBV has also been shown to increase the excitability of the corticospinal tract as assessed by motor evoked potential (MEP) amplitude in some muscles but not in others (Mileva et al., 2009; Krause et al., 2016; Pamukoff et al., 2016b, 2016a). Although several studies have investigated the effects of WBV on sensorimotor pathways, only one study has investigated how ULV affects sensorimotor transmission in the human upper limbs to date (Budini et al., 2017). It was determined that while neurologically intact participants held a vibrating handle, there was a decrease in forearm H-reflex amplitude that did not compromise manual dexterity or increase force fluctuations. Further investigation is required to effectively implement targeted rehabilitation training.

The overall objective of this study was to assess transmission along spinal and corticospinal pathways that control the human upper limb during ULV. Spinal pathways were assessed based on the amplitude of Hoffmann (H-) reflexes, cutaneous reflexes, and cutaneous conditioning of H-reflexes. Corticospinal pathways were assessed based on the amplitude of motor evoked potentials (MEPs) elicited using transcranial magnetic stimulation. Based on previous literature it was hypothesized that ULV would inhibit H-reflexes and facilitate MEPs recorded from the flexor carpi radialis (FCR) muscle compared to control trials with no vibration.

## 2 Methods

### Participants

Fourteen neurologically intact participants (10 male; 4 female, 29.4 ± 9.1 years, 174.2 ± 9.1 cm, 70.6 ± 11.8 kg) free from metabolic or neuromuscular disorders completed the experimental protocol which was approved by the University of Alberta Human Research Ethics Board. Participants were informed of all experimental procedures and signed a written consent form.

### Experimental procedure

The protocol directly compared two distinct tasks of ULV and CONTROL (no vibration) with the task order delivered in a random order between participants. The aim was to assess whether ULV altered sensorimotor transmission by assessing Hoffmann (H-) reflexes, cutaneous reflexes, and motor evoked potentials (MEPs) in the flexor carpi radialis (FCR) and biceps brachii (BB; MEPs only). For all outcome measures 20 stimuli were delivered (3-5 s apart) while participants held ≈10% of their maximum voluntary contraction (MVC) in FCR. For the trials that involved ULV, the vibration was delivered to the right upper limb using a custom-built vibration device that participants held with their right hand at all times during the experiment. Participants remained seated with their body and arm position maintained in a consistent position throughout the duration of the experiment using restraints (Figure 1). For all conditions, participants maintained a consistent contraction of ~10% of peak muscle activity of the right FCR using visual feedback of the rectified and low pass filtered EMG signal displayed on a computer screen. This was done to ensure similar excitability of the FCR spinal motor pool throughout all conditions. Elbow angle was maintained throughout the experiment within each participant between 100 - 110°. Rest periods were incorporated throughout to avoid fatigue.

**Figure 1.**
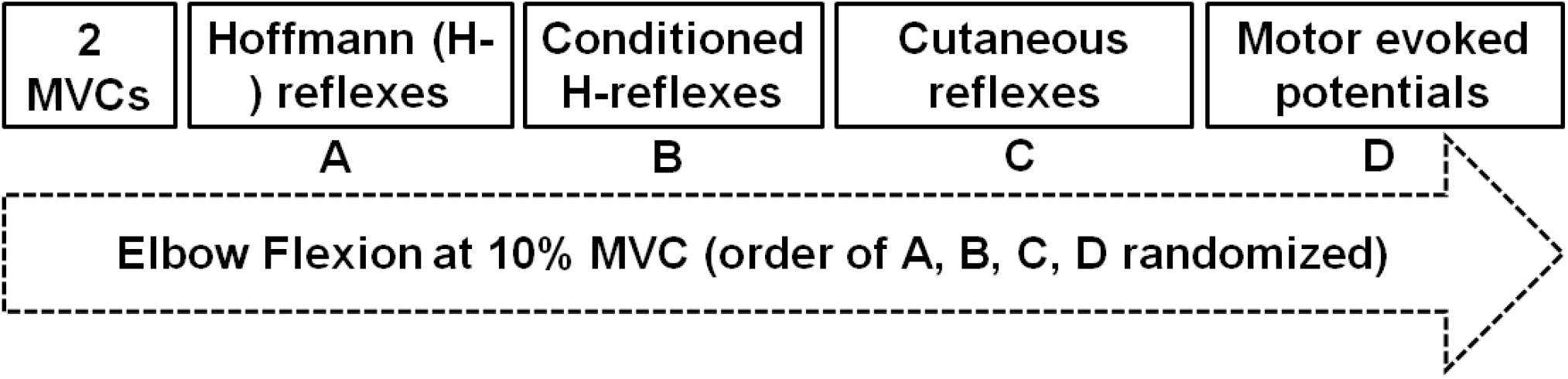
Experimental protocol. Two maximal voluntary contractions were followed by twenty evoked responses during ≈10% peak muscle activation of the flexor carpi radialis (FCR) in a randomised order of A) Hoffmann (H-) reflexes; B) Conditioned H-reflexes; C) Cutaneous reflexes; D) Motor evoked potentials.

### Upper limb vibration

A custom built vibration device was used to deliver ULV to the right upper limb through the hand while participants actively gripped the device (Pujari et al., 2016) (Figure 1). The ULV was maintained at a displacement amplitude of 0.353 mm, a frequency of 30 Hz and acceleration of 1.286 m/s^2^. A built-in load cell assessed force during isometric elbow and wrist flexion (Pujari, 2016; Pujari et al., 2019b). For trials involving ULV, the vibration was turned on and data collection started within ~ 1 min, remained on for the duration of the trial (~5min) and the vibration was turned off immediately after data were collected (>1 min).

### Electromyography (EMG)

Surface electromyography (EMG) was recorded through electrodes placed on the skin over the right flexor carpi radialis (FCR) and biceps brachii (BB) as shown in Figure 1A (2.25 cm2; Vermed Medical, VT, USA). The skin was cleaned with alcohol and then electrodes were placed in a bipolar configuration longitudinally along the predicted fibre direction in accordance with SENIAM procedures (Hermens et al. 2000). EMG signals were amplified 2000x, band pass filtered at 20Hz to 1000Hz (NeuroLog System; Digitimer, WelwynGarden City, UK) and then digitized at 2000 Hz (National Instruments Corp. TX, USA) using custom-written continuous acquisition software (LabVIEW, National Instruments, TX, USA).

### Maximal voluntary contractions

Three MVCs of the elbow and wrist flexion while gripping the ULV device were performed at the beginning of each experiment in the same position as all experimental conditions (Figure 1). Verbal encouragement and visual feedback were provided to ensure peak force and muscle activity were achieved. Each MVC was held for ≈3 seconds with one minute of rest provided between each attempt. The MVC that elicited the most torque was the MVC used to normalize all subsequent torque measurements. The MVC torque was quantified over a 0.3 s window centered on the peak during each MVC.

**Figure 2.**
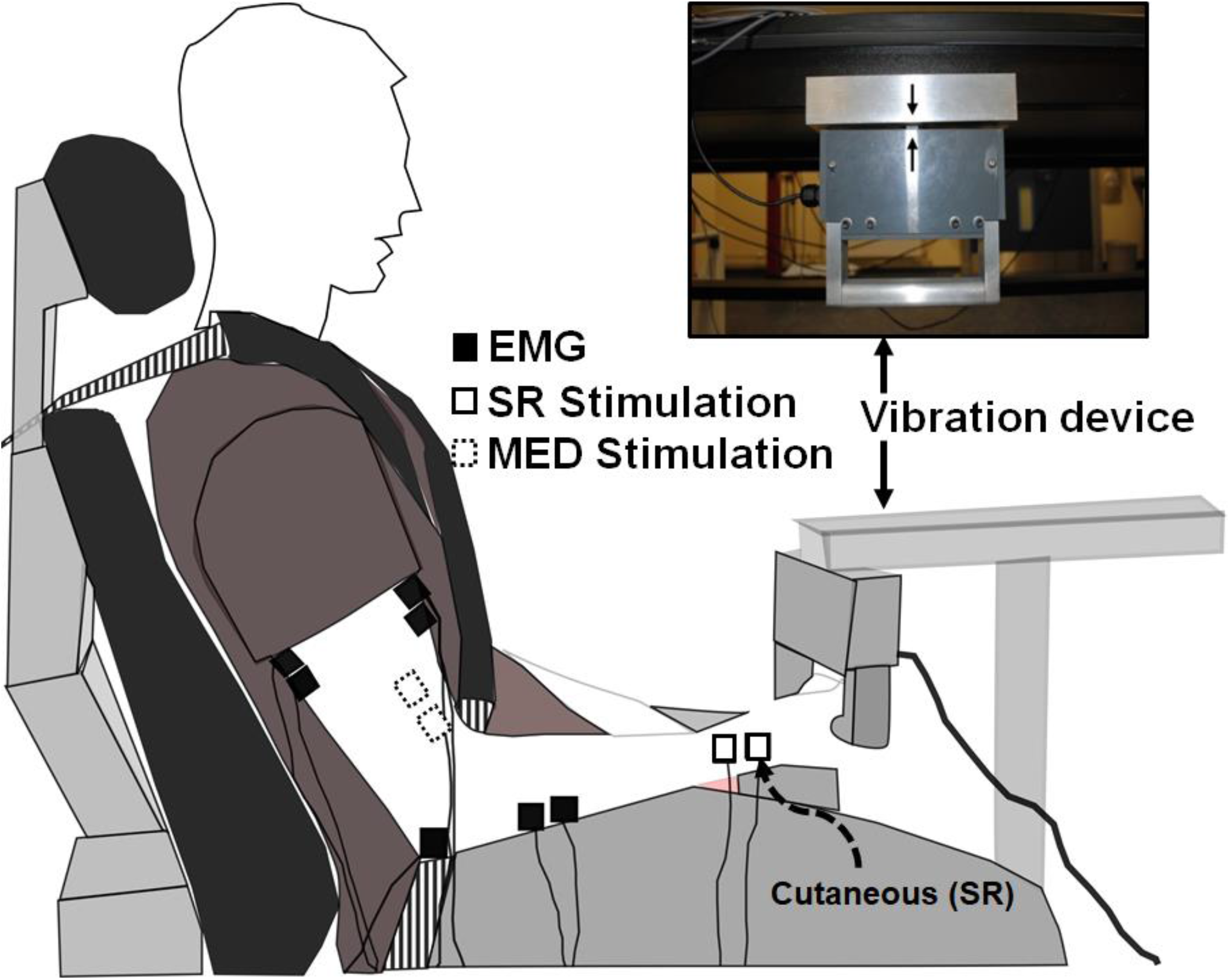
Experimental setup. White boxes with dashed lines indicate the stimulation electrode placement on the median nerve proximal to the elbow on the inside of the arm to elicit Hoffmann (H-) reflexes. White boxes with solid lines indicate the stimulation electrode placement on the superficial radial nerve at the wrist to both elicit cutaneous reflexes and condition H-reflexes during separate trials. Black boxes indicate recording electrode placement to determine H-reflexes, cutaneous reflexes, and MEPs with electromyography (EMG).

### H-reflexes

To evoke H-reflexes, 1 ms pulses were delivered through self-adhesive electrodes (Vermed Medical, VT, USA) over the median nerve proximal to the elbow using a Digitimer (DS7A or DS7AH) stimulator. At the beginning of each experiment stimulation intensity was adjusted to identify that which evoked H-reflexes that were accompanied by small M-waves, were on the ascending limb of the H-reflex recruitment curve and were ≈70% of the maximal H-reflex (H_max_). For each participant M-wave amplitude was maintained across all conditions by adjusting stimulation intensity in 1 mA increments as needed to ensure similar motor and sensory axons were recruited across conditions. Five maximal motor responses (M_max_) were also recorded by stimulating the median nerve at 1.25x the minimum intensity required to evoke M_max_. M_max_ was used to normalize H-reflexes, cutaneous reflexes, and MEPs.

### Cutaneous reflexes

Trains of stimuli (5 × 1ms @ 300 Hz) were applied to the superficial radial (SR) nerve just proximal to the radial head (Zehr and Hundza, 2005; Barss et al., 2020) were used to both condition H-reflexes and elicit cutaneous reflexes. A Grass S88 stimulator, SIU5 stimulus isolation and CCU1 constant current unit (Grass Instruments, Austin, TX, USA) (Nakajima et al., 2013) were used to deliver stimulation. Radiating thresholds (RT) to SR nerve stimulation were identified for each participant and were used to determine stimulation intensity. RT was defined as the lowest intensity that produced radiating paresthesia in the entire cutaneous receptive field of the SR nerve (Duysens et al., 1990; Brooke et al., 1997). Stimulation for each participant was set at 3xRT to evoke cutaneous reflexes.

### Somatosensory conditioning of H-reflexes

To explore potential presynaptic effects, a conditioning-test stimulation paradigm was incorporated that is known to reduce pre-synaptic inhibition in the FCR, facilitating H-reflex amplitude (Nakajima et al., 2013; Barss et al., 2018). SR stimulation was delivered at 2xRT 37 ms prior (conditioning-test interval) to proximal median nerve stimulation (H-reflex). Twenty pulses were applied during separate trials every 3 – 5 s at the intensity required to evoke the same amplitude M-wave as during the unconditioned reflexes.

**Figure 3.**
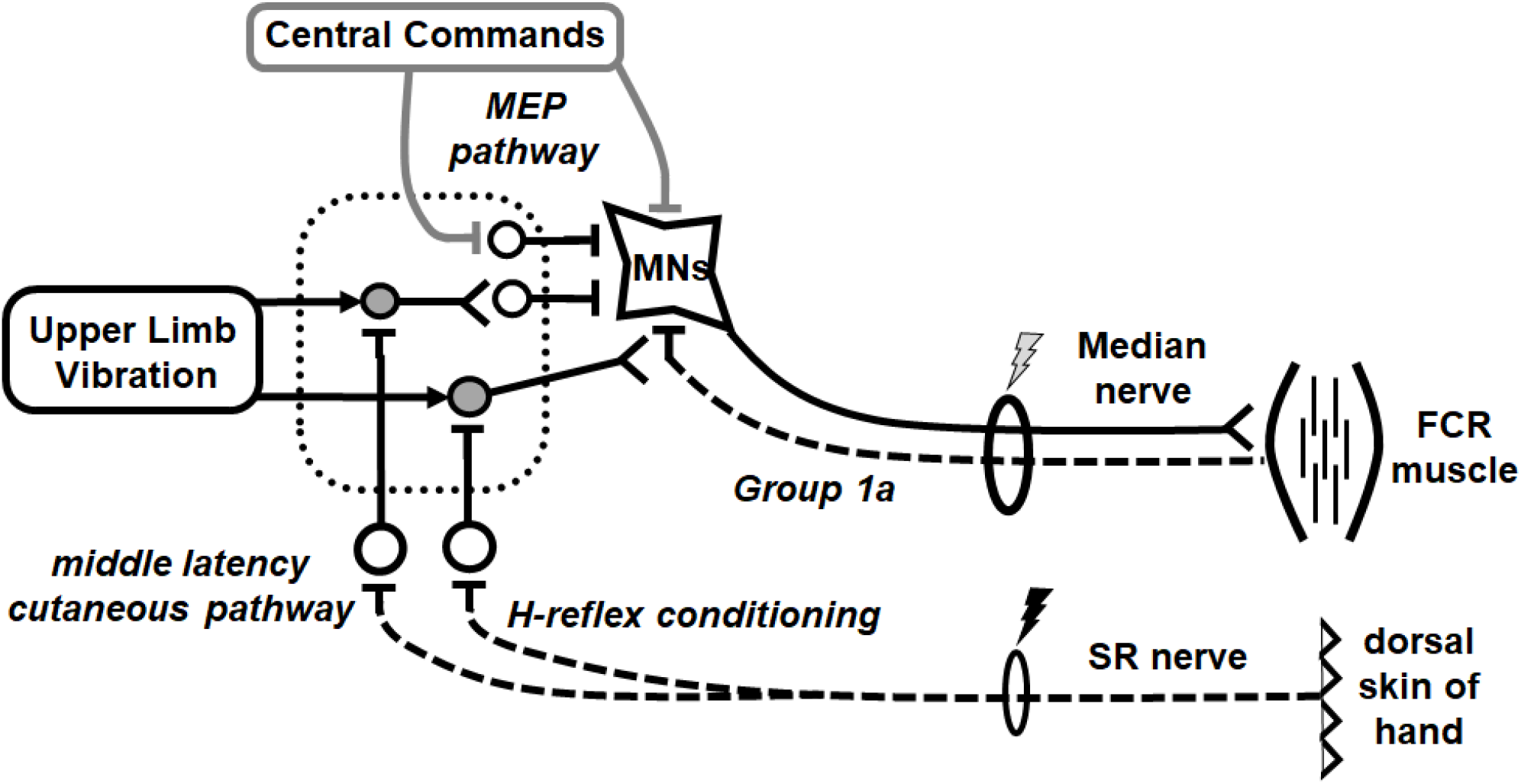
Schematic diagram outlining likely neural pathways for integration of inputs from indirect vibration applied to the upper limb (ULV). It remains likely that ULV has both pre- and post-synaptic effects on spinal excitability without altering cortico-spinal excitability. The mechanism of H-reflex conditioning with superficial radial (SR) nerve stimulation reducing pre-synaptic inhibition of the FCR Ia afferent is highlighted. Primary afferents are displayed with dashed lines. Excitatory synapses are displayed as a “**T**” with open cell bodies while inhibitory synapses are displayed with a “**V**” and grey cell bodies. The dotted rectangle represents a network of interneurons within the spinal cord.

### Transcranial magnetic stimulation

Transcranial magnetic stimulation (MagPro R30, Medtronic) was delivered over the left motor cortex to elicit motor evoked potentials (MEPs) in the FCR and BB muscle to test corticospinal excitability. The location for stimulation was chosen by first determining the optimal site for FCR MEPs by periodically moving the coil to identify the location that produced the largest MEP. This location was then marked and maintained within 1 mm relative to cortical landmark throughout the experiment using an image guidance system (Brainsight, Rogue Research) to ensure accurate and consistent stimulation. Stimulation intensity was set at the beginning of the experimental protocol and maintained across ULV and CONTROL conditions to evoke an MEP that was ≈70% of the maximal MEP amplitude so that both inhibition and facilitation of the MEP could occur.

### Data collection and analysis

FCR H-reflex, M-wave, cutaneous reflex, and MEP amplitudes were averaged across twenty sweeps for each condition and analysed offline using Matlab 2019© (Mathworks, Nantick, MA). The peak-to-peak amplitude were quantified for M-waves, H-reflexes and MEPs. Cutaneous reflexes were quantified by averaging twenty responses to SR stimulation then subtracting 50 ms of pre-stimulation muscle activity, leaving reflex activity to be assessed (Brooke et al., 1997; Zehr and Stein, 1999). Background muscle activity was quantified as the averaged EMG activity of a 50ms pre-stimulus window for each condition. The stimulus artifact was removed from the subtracted reflex trace and data were then low-pass filtered at 30 Hz using a dual-pass, fourth order Butterworth filter. The peak short (40-70 ms post-stimulus), middle (70-110 ms post-stimulus) and long latency responses (110-140 ms post-stiulus) were evaluated (Zehr et al., 1997, 1998)(Figure 5A). The time window for each latency was visually chosen around the peak response (Either excitation or inhibition). EMG was then averaged over a 10 ms window centered around the maximum response to obtain a single value.

### Statistical Analysis

Dependent measures of background muscle activity, M-wave, H-reflex, cutaneous reflex amplitudes, and MEPs were assessed using SPSS Statistic 20 (Chicago, IL). M-waves and H-reflexes were analyzed using a 2 (CONTROL vs ULV) x 2 (Unconditioned vs Conditioned) repeated measures analysis of variance (rmANOVA). Background muscle activity, MEPs and cutaneous reflexes were analyzed using paired sample t-tests (CONTROL vs ULV). When appropriate, post-hoc analyses were performed using Bonferroni corrected pairwise comparisons. Significance was accepted below *p*=0.05.

## 3 Results

### Background muscle activity

Background EMG activity in FCR was not different between ULV and CONTROL for all dependent measures (*p* > 0.05).

### M-waves, H-reflexes and conditioned H-reflexes

Representative examples of FCR M-waves and H-reflexes from one participant recorded during the unconditioned CONTROL and ULV trials are shown in Figure 4A. For this participant, when M-wave amplitudes were of similar amplitude, H-reflexes were smaller during ULV (vibration) than no-vibration (control) trials. A 2 × 2 repeated measures ANOVA confirmed that for the group M-wave amplitudes were not different between ULV and CONTROL tasks or between conditioned and unconditioned trials (Figure 4B), there was no significant interaction (F_(1, 13)_=1.819, *p* = 0.200) or main effects of vibration (F_(1, 13)_=0.096, *p* = 0.762) or conditioning (F_(1, 13)_=2.478, *p* = 0.139).

**Figure 4:**
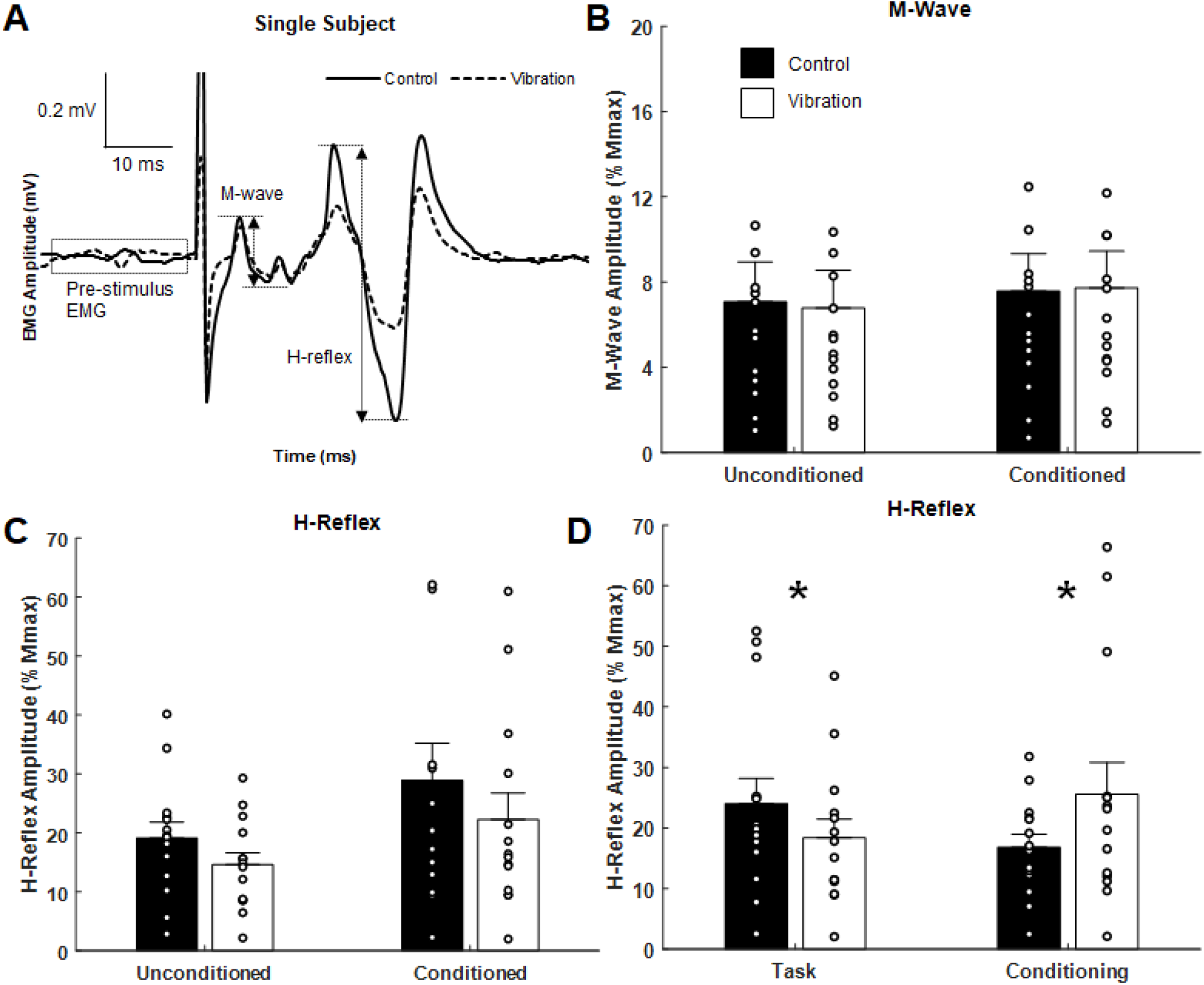
Effects of upper limb vibration on Hoffmann (H-) reflex. A) Single subject traces highlighting the suppression of H-reflex amplitude during ULV while M-wave is maintained constant. Solid traces indicate the average of 20 sweeps during CONTROL, whereas dotted traces indicate the average trace during ULV. B) Group average of M-wave amplitude across conditions indicating the same descending input was provided across condition. C) Group averages of H-reflex amplitude with and without ULV for both unconditioned and conditioned reflexes. D) Group average of H-reflex amplitude pooled across task (ULV vs CONTROL) and effect of conditioning. All single subject data is included as clear circles. * Indicates significant difference in H-reflex amplitude. Values are mean ± SD (*p* < 0.05).

A 2 × 2 repeated measures ANOVA indicated there was a significant main effect of both vibration (F_(1, 13)_ = 7.178, *p* = 0.019) and conditioning (F_(1, 13)_ = 5.124, *p* = 0.041) for H-reflex amplitude. Pooled across task, figure 4D highlights that SR nerve conditioning significantly facilitated H-reflex amplitude (25.6±5.2 % M_max_) compared to unconditioned reflexes (16.8±2.1 % M_max_). This highlights the conditioning paradigm was effective at reducing pre-synaptic inhibition and facilitating the H-reflex. Importantly, pooled across effect of conditioning, figure 4D indicates a significant reduction in H-reflex amplitude during ULV (18.4±3.1 % M_max_) compared to control (24.0±4.1 % M_max_). This signifies that ULV applied to the upper limb had a similar inhibitory effect on H-reflex transmission for both conditioned and unconditioned reflexes.

### Cutaneous reflexes

Representative examples of cutaneous reflexes recorded from one participant during CONTROL and ULV trials are shown in Figure 5A. Responses in the early, middle, and late latency components have been assessed separately. As shown in Figure 5C, a paired sample t-test indicated that there was significantly more inhibition of the middle latency response during ULV (−1.73 ± 2.2 % M_max_; F_(1, 13)_ = 9.279, *p* = 0.009) than CONTROL trials (−1.50 ± 2.1 % M_max_). There were no significant differences in the amplitudes of the early (1.4 ± 2.6 vs 1.5 ± 2.6 % M_max_; (F_(1, 13)_ = 0.307, *p* = 0.589) or late (0.94 ± 1.8 vs 0.98 ± 1.9 % M_max_; (F_(1, 13)_ = 0.077, *p* = 0.786) latency cutaneous responses (Figure 5 B/D).

**Figure 5.**
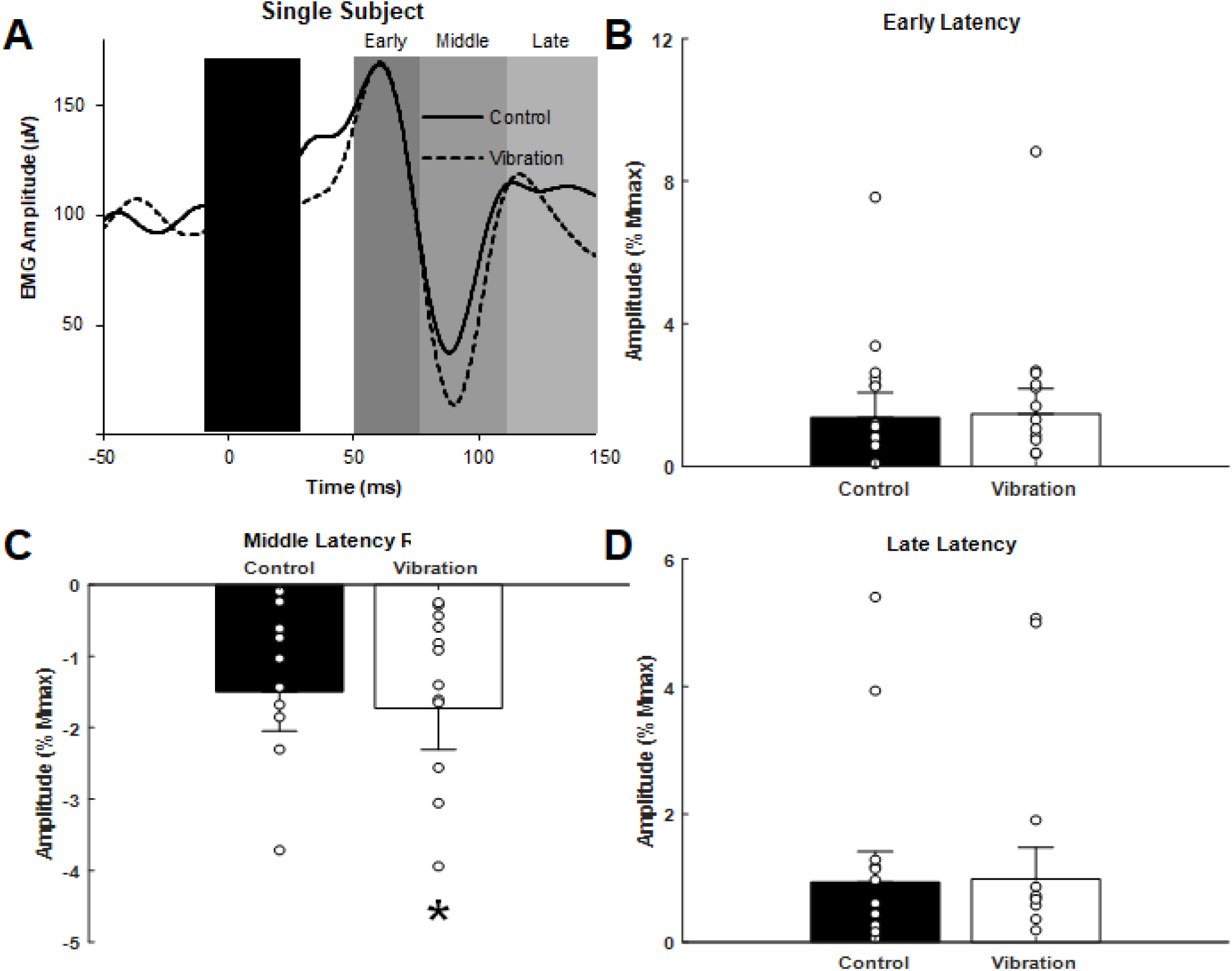
Effects of upper limb vibration on cutaneous reflex amplitude. A) Single subject traces providing representative examples of cutaneous reflexes. Solid traces indicate the average of 20 sweeps during CONTROL, whereas dotted traces indicate the average trace during ULV. B) Group average across conditions of early latency cutaneous reflex amplitude. C) Group average across conditions of middle latency cutaneous reflex amplitude. D) Group average across conditions of long latency cutaneous reflex amplitude. All single subject data is included as clear circles. * Indicates significant reduction in middle latency cutaneous reflex amplitude during ULV. Values are mean ± SD (*p* < 0.05).

### Motor evoked potentials

Representative examples of MEPs recorded from one participant for CONTROL and ULV are shown in Figure 6A. A paired sample t-test indicated there were no differences in MEPs between CONTROL and ULV in either the FCR (11.8 ± 8.2 vs 12.9 ± 9.7 % M_max_; (F_(1, 13)_ = 1.084, *p* = 0.317) or BB (4.8 ± 5.0 µV vs 4.7 ± 4.6 µV; (F_(1, 13)_ = 0.566, *p* = 0.465) (Figure 6B). This indicates that ULV did not alter corticospinal excitability for these muscles of the upper limb.

**Figure 6.**
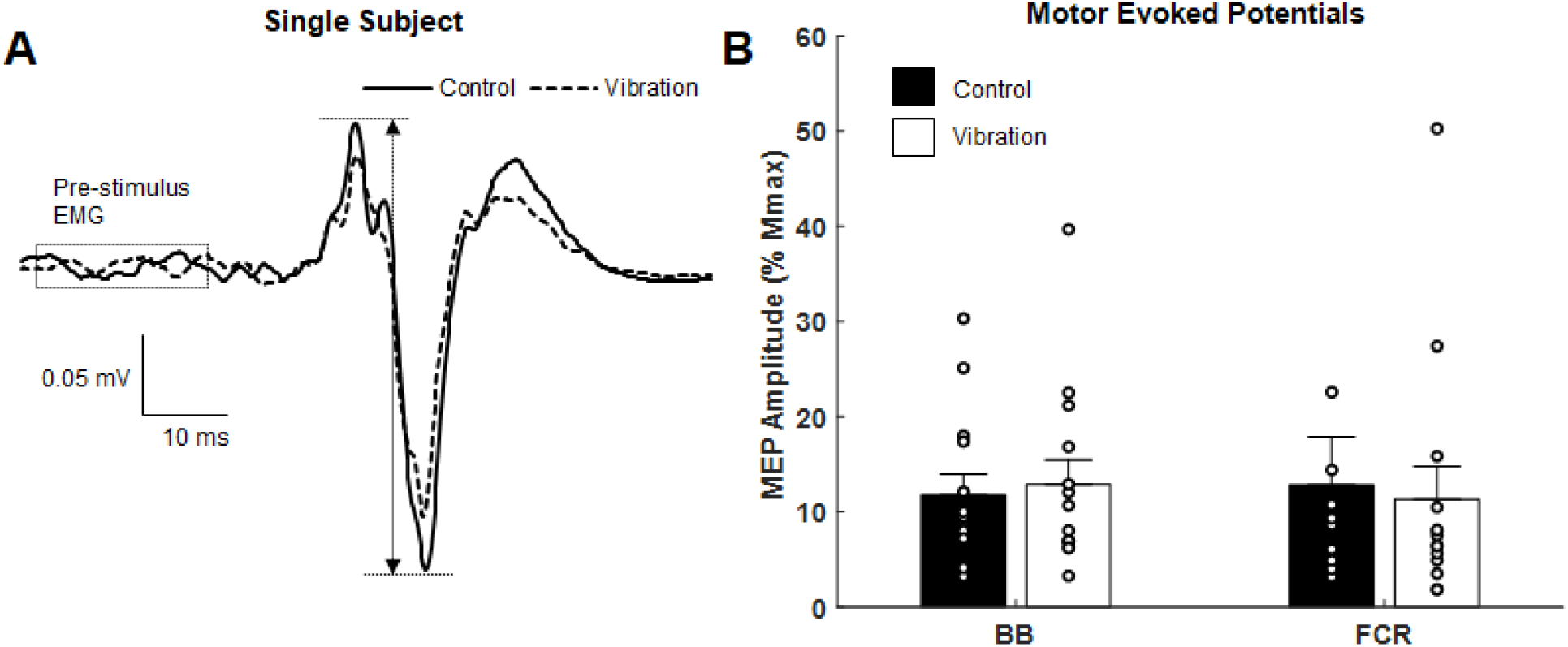
Effects of upper limb vibration on motor evoked potentials (MEPs). A) Single subject traces which show representative examples of MEPs during CONTROL and ULV. Solid traces indicate the average of 20 sweeps during CONTROL, whereas dotted traces indicate the average trace during ULV. B) Group average of motor evoked potentials during CONTROL and ULV in the flexor carpi radialis and biceps brachii muscles. All single subject data is included as clear circles.

## 4 Discussion

The study objective was to assess sensorimotor transmission in the human upper limb during ULV to identify potential corticospinal and spinally mediated sites of adaptation. Our hypothesis was supported in that ULV significantly inhibited H-reflex amplitude by 15.7% compared to CONTROL. ULV also strengthened the inhibition of middle latency cutaneous reflexes by 20.0% compared to control. Contrary to our hypothesis and previous WBV results, there was no significant effect of ULV on MEP amplitude in the upper limbs. This study highlights for the first time in the upper limbs that acute ULV inhibits spinally mediated neuronal circuits as demonstrated by the suppression of cutaneous and H-reflex responses without influencing corticospinal excitability of the forearm flexors. Together these results suggest that ULV increases pre-synaptic inhibition of afferent transmission.

### Upper limb vibration reduces H-reflex transmission

Previous investigations in the lower limbs of neurologically intact individuals have shown WBV significantly inhibits H-reflex amplitude (Armstrong et al., 2008; Kipp et al., 2011; Ritzmann et al., 2013; Hortobágyi et al., 2014; Ahmadi et al., 2015; Harwood et al., 2017; Laudani et al., 2018). H-reflex suppression has also been noted for individuals with a spinal cord injury although it was less pronounced compared to neurologically intact individuals (Sayenko et al., 2010). Results from the current investigation support both the previous literature (Budini et al., 2017) and the proposed hypothesis as ULV significantly inhibited FCR H-reflexes. Conditions were controlled to maintain consistent body position and descending voluntary drive between CONTROL and ULV conditions. Thus, the diminished H-reflex amplitudes during both ULV and WBV indicate and attenuation of spinal excitability of either a pre− or post-synaptic nature. This inhibition appears to be robust across both muscle group and source of indirect vibration. Figure 3 provides a schematic that highlights that ULV reduces H-reflex amplitude by either increasing pre-synaptic inhibition onto Ia afferents or providing an inhibitory post-synaptic input to the FCR motoneuron pool.

### Effectiveness of conditioning paradigm

A conditioning paradigm was employed to reduce pre-synaptic inhibition onto Ia afferents and facilitate the H-reflex (Nakajima et al., 2013) to determine potential pre-synaptic influences on H-reflex excitability during CONTROL and ULV. As expected, the conditioning paradigm was effective at facilitating excitability of the H-reflex pathway. The evoked motor response (M-wave) did not differ across conditioning paradigm or between CONTROL and ULV indicating the same relative input was provided into the spinal cord regardless of condition. With the same relative input, the conditioned H-reflexes depolarized more motor units likely due to a reduction in pre-synaptic inhibition, demonstrated by a significant increase in H-reflex amplitude (Nakajima et al., 2013). Although post-synaptic effects cannot be excluded in the current investigation since our conditioning stimulation was above the threshold of evoked responses in ongoing EMG.

Importantly, while the conditioning paradigm was effective at facilitating the H-reflex, applying ULV during a static task corresponds with an overall reduction in H-reflex excitability regardless of whether the reflex was conditioned. Therefore, it appears that an interaction occurred between the known conditioning input (Nakajima et al., 2013) and an inhibitory input from ULV which could be of either a pre- or post-synaptic nature. Figure 3 provides a schematic representation of the conditioning paradigm and these potential sites of interaction.

### Upper limb vibration strengthens inhibition of middle latency cutaneous reflexes

Applying ULV strengthened the inhibition of middle latency cutaneous reflexes. Cutaneous reflexes provide information on the relative contribution of sensory information from the skin being incorporated into ongoing motor output (Zehr and Stein, 1999). The convergence of excitatory and inhibitory effects on FCR motoneurons depends on the nerve being stimulated and the latency at which the response is measured. Similar to previous studies, middle latency responses to SR nerve stimulation in the FCR produce a large inhibitory effect (Zehr et al., 2001). Interestingly, when ULV was applied, there was significantly more inhibition of the middle latency response. Contributions to the inhibition of ongoing muscle activity at this latency likely occur at multiple levels of the spinal cord through converging pathways on the FCR motoneuron pool (Iles, 1996; Birmingham et al., 1998; Aimonetti et al., 1999; Zehr and Stein, 1999). Since the effect was not observed in either early or late latency reflexes during ULV, it is likely that the effects are occurring pre-synaptic to the motoneuron pool. The potential pre- and post-synaptic influences of ULV on middle latency reflex amplitude are shown in Figure 3.

### Upper limb vibration does not alter corticospinal excitability

Previous investigations have shown WBV increases corticospinal excitability of the tibialis anterior muscle while the vibration was being applied (Mileva et al., 2009), in the soleus muscle for up to 10 minutes after the application of vibration (Krause et al., 2016), and to the vastus medialis for up to 20 minutes (Pamukoff et al., 2016b). However, there were no significant differences in MEP amplitude of the gastrocnemius (Krause et al., 2016) or soleus (Mileva et al., 2009) within these same investigations. Also, no significant increases in MEP amplitudes occurred after WBV in the vastus medialis after ACL reconstruction (Pamukoff et al., 2016a). Thus, the reported increases in corticospinal excitability during and after WBV are not consistent across muscle groups or time points (during and after WBV). To our knowledge no studies exists on the effects of ULV on MEP amplitude. Contrary to our hypothesis, ULV did not alter corticospinal excitability for either the FCR or BB. This indicates that ULV did not alter the excitability of the motor cortex or the motoneuron pool. When the results are combined it suggests that ULV inhibits spinal reflexes primarily due to pre-synaptic mechanisms as shown in Figure 3.

### Functional implications

A specific target population for the incorporation of ULV are individuals with a spasticity-related deficiency in sensorimotor control which include spinal cord injury (DeForest et al., 2020), stroke (Liepert and Binder, 2010) and spastic movement disorders (Dietz and Sinkjaer, 2007). Results of this study indicate that ULV increases inhibition on spinal pathways that is likely of a pre-synaptic nature. Thus, ULV may be an effective way to reduce spasticity within a session of rehabilitation, in a similar fashion to direct vibration. Recently, direct vibration applied to either the Achilles or tibialis anterior tendon after spinal cord injury suppressed late spasm-like activity in antagonist but not agonist muscles, likely via reciprocal inhibitory mechanisms (DeForest et al., 2020). Thus, improving acute functional ability through ULV for individuals with neurological movement disorders may be effective strategy for targeted rehabilitation (Ahlborg et al., 2006; Ness and Field-fote, 2009; Sayenko et al., 2010). This provides the mechanistic basis for the targeted use of ULV to potentially reduce spasticity and improve rehabilitation outcomes.

### Limitations

It remains possible that due to technical limitations of the ULV apparatus used within the current investigation, the amplitude of vibration provided to the upper limb was not sufficient to alter corticospinal excitability. The maximal displacement amplitude of the ULV device used in the current investigation is 0.353 mm. In the lower limbs it has been shown that the electromyography response induced by WBV depends on vibration amplitude, frequency and muscle stretch; with vibration amplitudes tested ranging from 0.5mm to 1.5mm (Pujari, Amit N. et al., 2019). If a larger amplitude was employed, it remains possible that a greater suppression of cutaneous and H-reflexes would have occurred while altering corticospinal excitability as seen in previous investigations using WBV. Further exploration will be required to determine if the lack of MEP facilitation is due to the amplitude of vibration provided. It also remains possible that WBV may have more influence on corticospinal excitability during conditions of postural control and weight bearing compared to ULV. It will be important for future investigations to determine whether effects of ULV are muscle specific (flexor vs extensor; upper vs lower), task specific (standing vs grasping) and amplitude dependent to ensure it is implemented in the optimal contexts (Pujari, Amit N. et al., 2019).

## 5 Conclusion

A single session of ULV altered transmission along spinal but not corticospinal pathways as demonstrated by a significant inhibition of both Hoffmann (H-) and middle latency cutaneous reflex amplitudes while motor evoked potentials remained unchanged. Therefore, it appears acute ULV alters segmental sensorimotor transmission within the forearm flexors likely due to increased pre-synaptic inhibition. ULV may provide an effective avenue for targeted rehabilitation training after damage to the nervous system. Specifically, ULV may be used as a concomitant therapy to reduce acute spasticity allowing for enhanced effectiveness both within a rehabilitation session and over time.

## 7 Author Contributions

Experiments were conducted in the Human Neurophysiology Laboratory, University of Alberta, Canada. TSB, DFC and ANP conceptualised the study and experimental design. TSB, DM and ANP collected the data. TSB and ANP analysed the data. TSB, DFC and ANP contributed to the interpretation and drafting of the manuscript. TSB, DFC, DM and ANP provided revisions for and approved the final draft of the manuscript.

## 9 Abbreviations

H-reflex: Hoffmann reflex
M-wave: direct motor response
Hmax: maximally evoked H-reflex
Mmax: maximally evoked motor response
BB: Biceps Brachii
TB: Triceps Brachii
FCR: flexor carpi radialis
ECR: extensor carpi radialis
SR: superficial radial
MED: distal median
PT: perceptual threshold
RT: radiating threshold
CONTROL: no vibration stimulation
ULV: upper limb vibration

## 10 Acknowledgments

The authors wish to acknowledge the participants for their contributions during data acquisition.

